# Saturated mutagenesis screen of M-MLV reverse transcriptase identifies variants enhancing prime editing efficiency

**DOI:** 10.64898/2026.07.06.736660

**Authors:** Hui Li, Yifan Wang, Chenxin Zhang, Thi Tun Thi, Yu Shi, Kai Yan Cheng, Chunyi Hu, Haojie Yu

**Affiliations:** Precision Medicine Research Programme, Yong Loo Lin School of Medicine, National University of Singapore, Singapore; Cardiovascular Disease Research Programme, Yong Loo Lin School of Medicine, National University of Singapore, Singapore; Department of Biochemistry, Yong Loo Lin School of Medicine, National University of Singapore, Singapore; Department of Biological Sciences, Faculty of Science, National University of Singapore, Singapore 117543, Singapore

**Author notes:** Correspondence (H.J.Y.). Co-first author (Hui Li, Yifan Wang).

## Abstract

Prime editing enables the precise modification of genomes, thereby holding great potential for the treatment of genetic diseases. Despite substantial advancements in prime editing technology and the initiation of the first clinical trial for treating chronic granulomatous disease, further enhancement of editing efficiency across edit types is still urgently needed. Here, we developed a compact prime editor, PE2ΔR, by deleting the RNase H domain of the MMLV reverse transcriptase (MMLV-RT). We then conducted a saturated mutagenesis screen targeting two DNA interacting regions within the PE2ΔR-RT Fingers domain. By integrating three highly effective mutations (I61R, V101R, S67W) into PEmax lacking RNase H domain (termed PEmaxΔRM3), we achieved up to a 90% increase in editing efficiency across editing types compared to PEmax. Structural modelling using AlphaFold 3 suggests that these mutations enhance primer–template stabilization and guide the RNA/DNA hybrid into a catalytically favourable trajectory, providing a mechanistic explanation for the enhanced activity. Taken together, our study demonstrates proof-of-concept for the application of unbiased mutagenesis screen to identify novel mutations that enhance prime editor performance. Furthermore, we discovered that RT variants (I61R, V101R, S67W) synergize with PEmax and epegRNA to improve prime editing efficiency across edit types, with the strongest improvement observed in introducing small deletions.

## Background

Prime editing (PE) is a versatile genome-editing technology that enables precise modifications, including base substitutions, insertions, and deletions, without requiring double-strand breaks or exogenous donor DNA templates^1–3^. The prime editor enzyme (PE2) consists of an SpCas9 nickase (H840A) fused to the Moloney murine leukemia virus reverse transcriptase (MMLV-RT)^1–4^. This system is directed to a specific genomic site by a prime editing guide RNA (pegRNA), which carries both a spacer sequence for target recognition and a 3′ extension that serves as a template for new DNA synthesis ^1–3^. Upon target binding, Cas9 nickase introduces a single-strand break, exposing a 3′ DNA end that hybridizes with the 3′ extension of the pegRNA. The MMLV-RT then uses this sequence as a template to introduce the programmed edit, after which the newly synthesized DNA strand competes with the original strand for genomic incorporation ^1–3^.

Despite its broad potential, prime editing remains constrained by limited editing efficiency, which varies across different edit types and target loci. Numerous strategies have been explored to improve efficiency, including improving fusion protein expression levels^5^ , activity^6–14^, nucleus localization^15^, and pegRNA stability^16,17^. Despite these efforts, editing efficiency remains a significant bottleneck. In this study, we performed a high-throughput site-saturation mutagenesis screen targeting two DNA interacting polypeptide stretches of MMLV-RT. This approach led to the identification of novel variants that substantially enhanced prime editing efficiency.

## Results

### Development of a compact PE lacking the RNase H domain (PE2ΔR)

Previous studies have demonstrated that M-MLV RT lacking RNase H activity exhibits enhanced cDNA synthesis efficiency and thermostability in the presence of a template- primer complex, likely due to increased DNA:RNA complex stability^4,18^. To investigate whether deletion of the RNase H domain affects PE editing efficiency, we constructed a truncated PE variant (PE2ΔR) by removing the RNase H domain (residues L499 to P677) (Supplementary Sequence 1). This truncated RT, encoded by 1,494bp, is 26.4% smaller than the wild-type MMLV-RT (Figure 1a).

**Figure 1.**
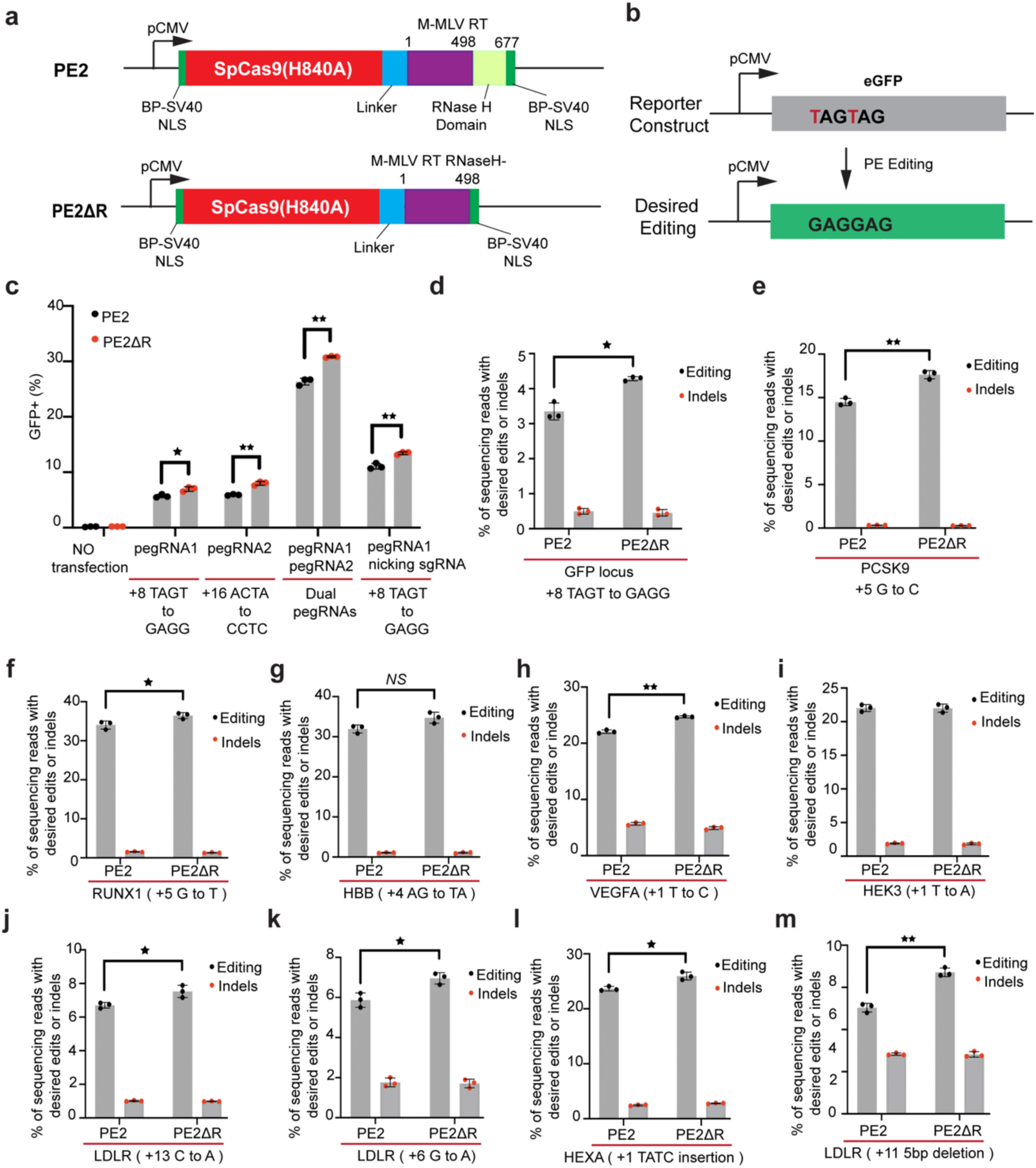
Deletion of RNase H domain in M-MLV RT enhances prime editing efficiency. **a.** Schematic representation of PE2 and PE2ΔR, highlighting the deletion of RNase H domain. **b.** Diagram illustrating the TAGTAG to GAGGAG transition, which converts two stop codons to Glu-Glu thereby restoring the function to a eGFP reporter in HEK293T cells. **c.** Frequencies of eGFP+ cells resulting from desired editing by PE2 and PE2ΔR were quantified by flow cytometry. **d.** Comparison of editing efficiency in eGFP reporter cells via NGS analysis. Indels represent unintended mutations that do not produce the desired sequence change. **e-m.** Comparison of editing efficiency between PE2 and PE2ΔR for nucleotide substitution at *PCSK9* locus **(e)**, *RUNX1* locus **(f)**, *HBB* locus **(g)**, *VEGFA* locus **(h)**, *HEK3* locus **(i)**, *LDLR* exon 4 **(j)**, and *LDLR* exon 10 **(k)**; for a 4-bp insertion at *HEXA* locus **(l)**; and for a 5-bp deletion at *LDLR* locus **(m)**. Results are from three independent experiments. * p< 0.05, ** p< 0.01 by Student’s *t-*test between PE2 and PE2ΔR. Error bar in c-m indicate mean ± SD.

We next evaluated the editing efficiency of PE2 and PE2ΔR using HEK293T eGFP reporter cells containing two premature TAG stop codons that block functional eGFP expression (Figure 1b and Supplementary Sequence 2). Both PE2 and PE2ΔR were directed by two different pegRNAs (targeting sites upstream or downstream of the editing site) designed to convert TAGTAG to GAGGAG, with or without a nicking sgRNA. In addition, we explored whether simultaneous use of dual pegRNAs- encoding the same edits for the forward and reverse DNA strands- could further enhance editing efficiency^19^. Flow cytometry analysis revealed that PE2ΔR achieved a 1.2-1.3 fold increase in editing efficiency compared to PE2 (Figure 1c and Figure s1a). This improvement was further validated by next-generation sequencing (NGS), which confirmed significantly higher editing efficiency with PE2ΔR (Figure 1d). To determine whether further truncation of the C-terminus of RTΔR would affect editing efficiency, we generated a series of truncation variants by deleting up to 25 amino acids from the C-terminus of RTΔR. We observed that deletions exceeding 10 amino acids significantly compromised the enhanced editing efficiency of RTΔR (Figure s1b and s1c).

To further characterize the activity of the PE2ΔR, we assessed its performance in nucleotide conversions at eight endogenous loci using pegRNAs or epegRNAs (Figure 1e to 1l). Consistent with previous report^12^, PE2ΔR outperformed the full-length MMLV-RT at six out of eight loci in HEK293T cells (Figure 1e to 1l). Moreover, PE2ΔR demonstrated superior efficiency in introducing small deletions compared to PE2 (Figure 1m).

### High-throughput saturated mutagenesis screen identifies novel RT variants capable of enhancing prime editing efficiency

Previous studies suggest that prime editing efficiency is limited by the interaction strength between the PE-pegRNA complex and the target locus^5,20,21^. It has been reported that mutations in reverse transcriptase (RT) that enhance its interaction strength with the DNA:RNA hybrid, improve thermostability, or increase processivity are likely to improve prime editing efficiency^1,4,6^. During reverse transcription mediated by MMLV-RT, specific regions within the fingers domain- S60-Q84 and N95-D124- have been shown to directly interacts with the DNA template (Figure 2a and Supplementary Sequence 3)^4,22^. Based on these findings, we hypothesized that engineering these two DNA-binding stretches within the fingers domain could enhance the RT-DNA:RNA interaction strength and stability, thereby improving the prime editing efficiency.

**Figure 2.**
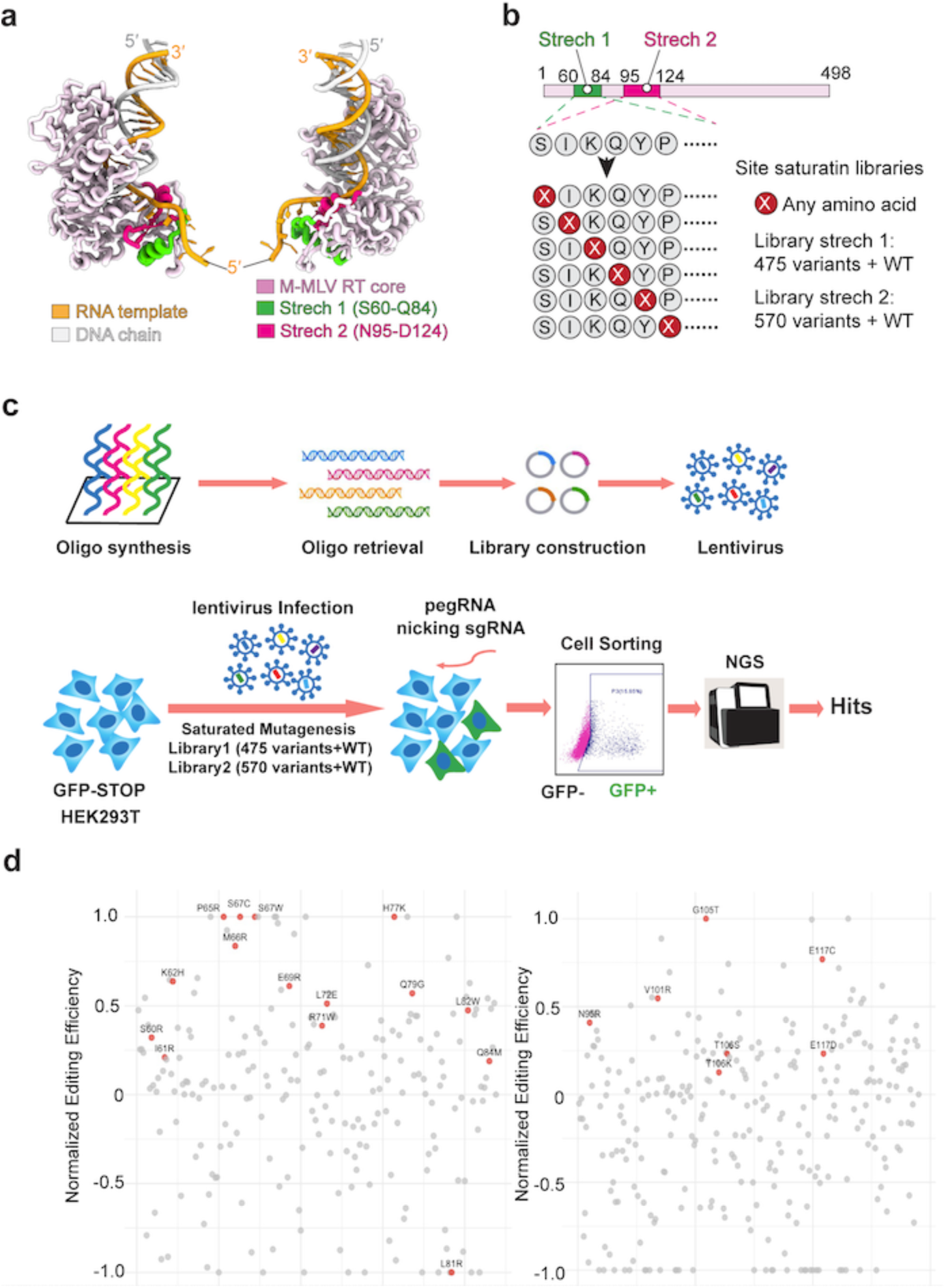
Saturated mutagenesis screening of RT DNA-interacting motifs identifies novel variants that enhance prime editing efficiency. **a.** Two orthogonal views of the M-MLV RT catalytic core (light pink) bound to an RNA/DNA hybrid (RNA template – orange; DNA product – grey). Two surface stretches chosen for engineering are highlighted: Stretch 1 spanning residues S60–Q84 (green) and Stretch 2 spanning residues N95–D124 (magenta). **b.** Schematic of the saturation-mutagenesis design. A linear map of the 498-residue RT marks Stretch 1 and Stretch 2 (green and magenta boxes). Every position within each stretch (red X) was diversified to any amino acid, producing Library Stretch 1 (475 variants + WT) and Library Stretch 2 (570 variants + WT).**c.** Schematic diagram showing the lentivirus-based mutagenesis screening in eGFP reporter cells. **d.** Scatter plots of variant enrichment from library 1 (left) and library 2 (right), showing normalized deep sequencing read counts relative to the wild-type control. Briefly, log2((abundance of variant in GFP positive cells)/(abundance of variant in GFP negative cells)) was calculated for each variant and for the wild-type construct. The editing efficiency of all variants were then normalized and scaled between -1 to 1, with the wild-type sequence set as the baseline (effect=0).

To identify novel mutations that enhance editing efficiency in a high-throughput and unbiased manner, we employed highly parallel oligonucleotide synthesis to construct two lentivirus-based PE2ΔR libraries L1 and L2 (Table S5). These libraries encoded all possible single-amino acid substitutions for the S60-Q84 and N95-D124 regions within the fingers domain, respectively, totalling 475 and 570 variants (Figure 2b and Figure S2). To enable screening, we generated an isogenic eGFP reporter cell line by isolating a single colony from bulk lenti-eGFP transduced cells. These reporter cells were then transduced with each lentiviral library, followed by antibiotics selection and expansion. After editing using a PE3 approach, we performed flow cytometric sorting to isolate eGFP+ (successfully edited) and eGFP-(unsuccessfully edited) cell populations. NGS analysis was conducted on these two populations, as well as on a whole library control, to identify RT variants exhibiting higher enrichment in the eGFP+ population compared to the parental RT (Figure 2c). Among all 1045 variants screened, sequencing analysis revealed that 176 variants show increased editing efficiency compared to parental RT (Figure 2d, Figure S3, Table S6 and S7).

### Functional validation of novel RT variants across edit types and genomic loci

We subsequently conducted functional validation of twenty-two prioritized variants hypothesized to enhance prime editing efficiency, as well as one variant hypothesized to reduce editing efficiency. These variants were tested in the eGFP reporter cell line by measuring the percentage of eGFP positive cells following successful editing (Figure 3a). In line with the screening result, PE2ΔR-L81R substantially reduced the editing efficiency compared to PE2ΔR (Figure 3a). Among the other 21 individual variants tested, the editing efficiency was enhanced by up to 59% compared to PE2, indicating that our saturated mutagenesis screen successfully discovered novel variants that enhance prime editor efficiency (Figure 3a).

**Figure 3.**
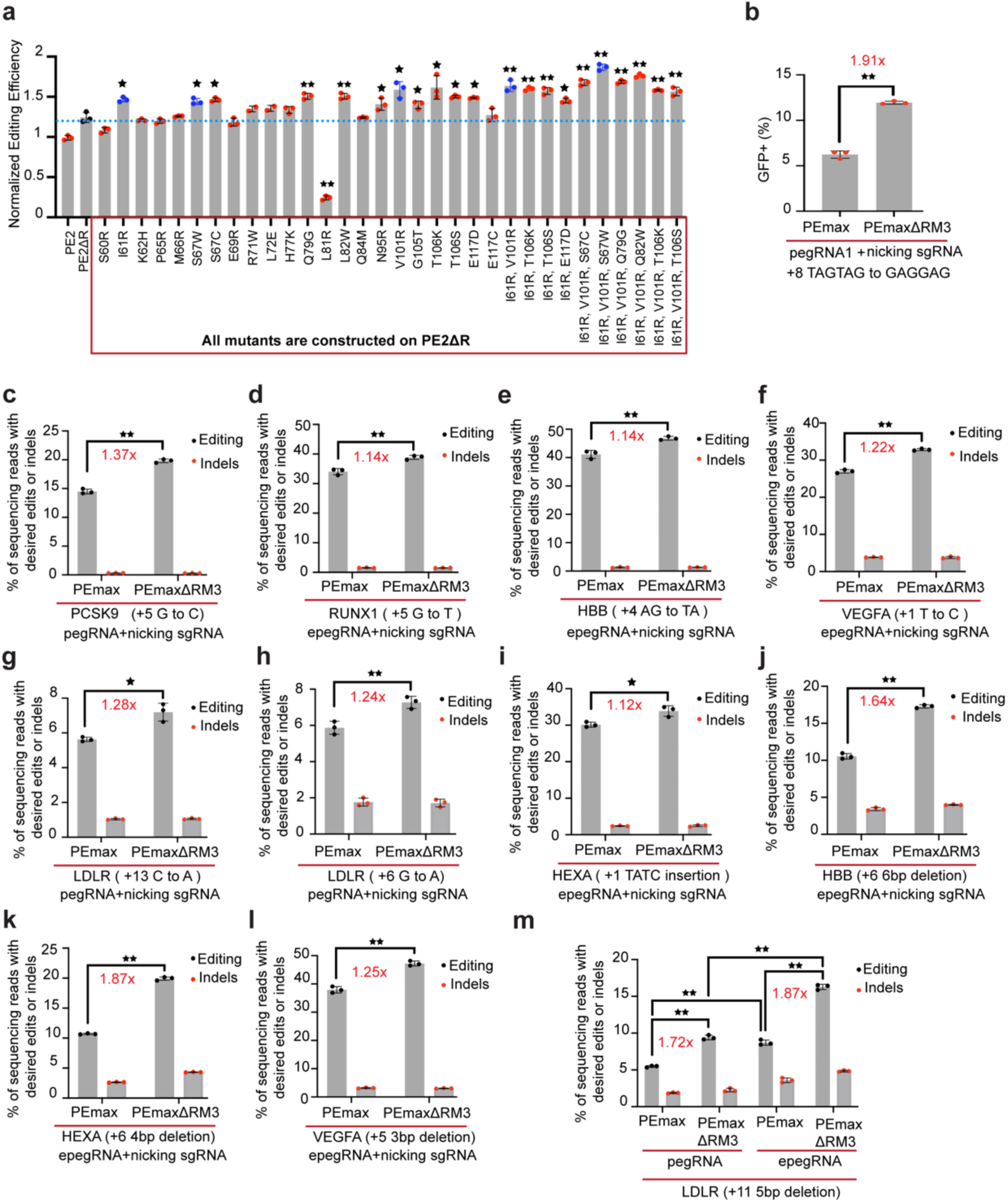
PEmaxΔRM3 increases editing efficiency across various endogenous loci and editing types. **a.** Normalized editing efficiency of PE2ΔR and its single or combined mutations in eGFP reporter cells. The percentage of eGFP+ cells after prime editing was measured by flow cytometry and subsequently normalized to PE2. **b.** Frequencies of eGFP+ cells resulted from desired editing by PEmax and PEmaxΔRM3 were quantified by flow cytometry. **c-i.** Comparison of editing efficiency between PEmax and PEmaxΔRM3 for nucleotide substitution at *PCSK9* locus **(c)**, *RUNX1* locus **(d)**, *HBB* locus **(e)**, *VEGFA* locus **(f)**, *LDLR* exon4 **(g)**, *LDLR* exon 10 **(h)**; for a 4-bp insertion at *HEXA* locus **(i)**. **j-m**. Comparison of editing efficiency between PEmax and PEmaxΔRM3 for small deletion at *HBB* locus **(j)**, *HEXA* locus **(k)**, *VEGFA* locus **(l)**, and *LDLR* locus using pegRNA or epegRNA **(m)**. Results were obtained from three independent experiments. * p< 0.05, ** p< 0.01 by Student’s *t-*test. Error bar in a-m indicate mean ± SD.

To explore whether combining these variants could further improve performance, we tested several combinations in eGFP reporter cells. Notably, the triple mutant PE2ΔR-I61R, V101R, S67W (referred to as PE2ΔRM3, Supplementary Sequence 4) exhibited a remarkable 86% increase in editing efficiency compared to the parental PE2 (Figure 3a). Specifically, the percentage of eGFP+ cells rose from 9.6% with PE2 to 17.9% with PE2ΔRM3 (Figure 3a). This synergistic effect highlights the potential of combining beneficial mutations to further optimize prime editing systems.

We next evaluated whether the three mutations (I61R, V101R, S67W) identified in PE2ΔRM3 could similarly enhance editing efficiency in an optimized SpCas9-based prime editor, PEmax^7^, in eGFP reporter cells. Consistent with the results from PE2ΔRM3, the corresponding PEmax variant (PEmaxΔRM3, which includes the deletion of RNase H domain and the introduction of I61R, V101R, S67W) increased the editing efficiency by 91% compared to the parental PEmax (Figure 3b, Figure S4, and Supplementary Sequence 5). This demonstrates that the beneficial effects of these mutations are not limited to a specific prime editor architecture but can be generalized to other optimized systems.

To further validate the utility of PEmaxΔRM3, we assessed its performance across a broader range of editing contexts. Using twelve pegRNA/nicking sgRNA pairs targeting seven endogenous genomic loci, we evaluated the efficiency of base substitutions, small insertions, and small deletions. For base substitutions and small insertions, PEmaxΔRM3 increased the desired editing efficiency by 12%-37% compared to PEmax (Figure 3c-3i). Notably, for small deletions, PEmaxΔRM3 achieved up to a 87% improvement in editing efficiency across five pegRNA/nicking sgRNA pairs (Figure 3j to 3m). Additionally, we observed that PEmaxΔRM3 synergizes effectively with engineered pegRNAs (epegRNAs)^17^. When paired with epegRNAs, PEmaxΔRM3 consistently outperformed the corresponding pegRNAs, underscoring the compatibility of these RT variants with advanced pegRNA designs (Figure 3m).

### Structural Modelling Reveals Mechanism Underlying Enhanced Activity of PEmaxΔRM3

To understand the molecular basis underlying this enhanced activity, we turned to structural modelling using AlphaFold 3. Structural analysis illuminates the synergy of this triple mutant. In the Alphafold 3 model of the RT–RNA/DNA complex (Fig. 4a), all three residues cluster immediately 5′ to the RNA template; this position is critical for shepherding the nascent hybrid into the polymerase active site. Close-up inspection (Fig. 4b) reveals that replacing Ile-61 and Val-101 with arginine introduces two basic side chains that form salt bridges with successive phosphates at positions –2 and –3 of the template, pinning the RNA in a catalytically productive trajectory. The S67W mutation, situated one helical turn upstream, stacks its indole ring against the ribose/base stack at position –1, further dampening template fraying. Together, these contacts rationalize the elevated cDNA synthesis we observe in vitro.

**Figure 4.**
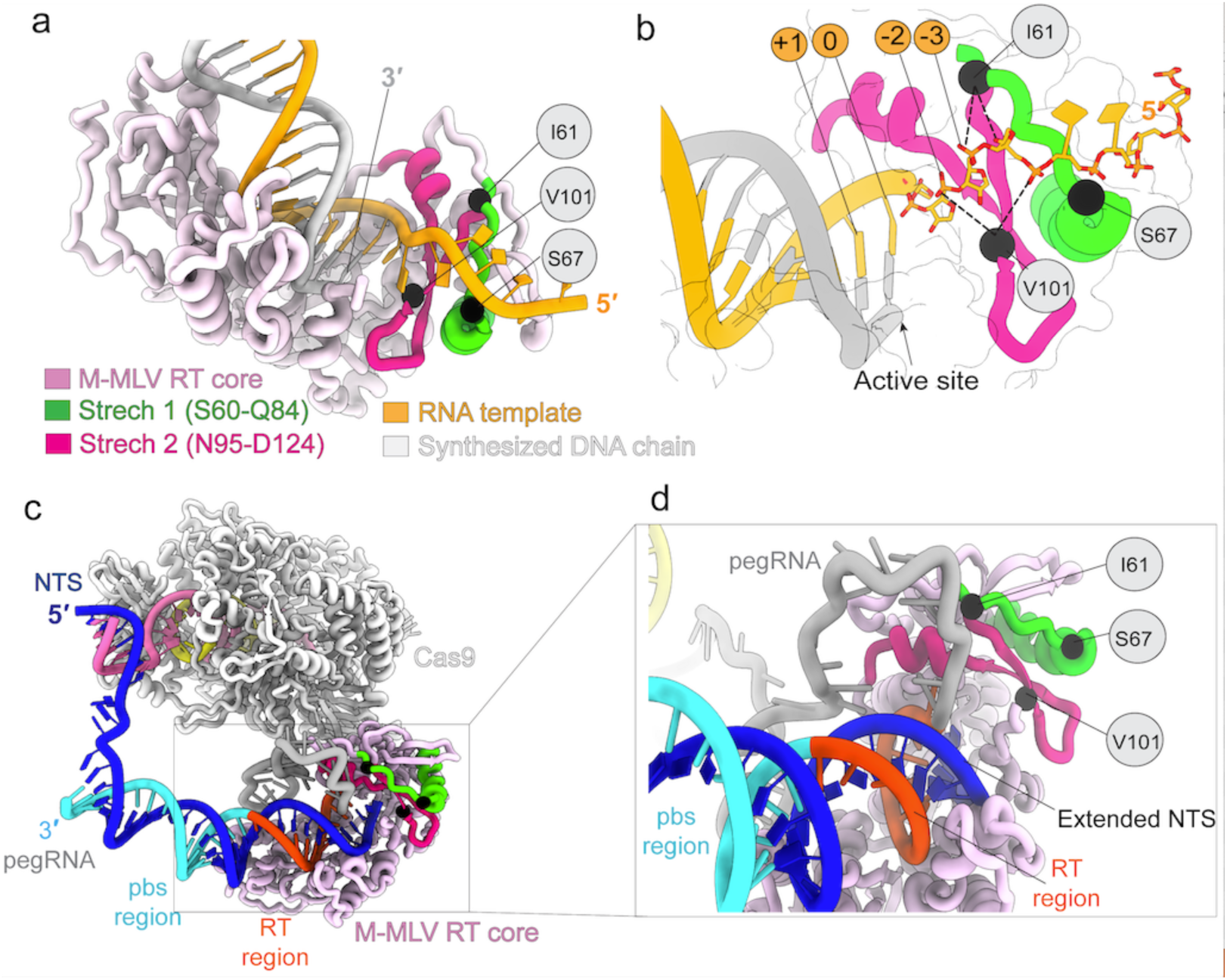
Structural basis for the activity-enhancing I61R/S67W/V101R triad in PEmaxΔRM3. **a.** Alphafold3 structure of the M-MLV reverse-transcriptase (RT) core (light pink) bound to an RNA/DNA primer–template duplex (RNA, orange; nascent cDNA, grey). Black spheres mark the three gain-of-function residues (I61, S67 and V101) whose substitutions define PEmaxΔRM3. **b.** Close-up of the template-entry channel. Template nucleotides are numbered relative to the polymerase catalytic centre (0). The I61R and V101R arginine side chains (modelled) project toward backbone phosphates at positions –3 and –2, whereas S67W contributes an indole π-stack beneath the –1 nucleotide. Together these contacts form a contiguous electrostatic/π-stacking clamp that stabilises the downstream RNA leader. **c.** AlphaFold 3 multimer model of the complete prime-editing complex after strand nicking. SpCas9 (grey) grips the target DNA duplex; the pegRNA primer-binding site (PBS, cyan) is annealed to the nicked non-target strand (NTS, dark blue), while the RT template portion of the pegRNA (orange) threads into the engineered RT core (light pink). **d.** Magnified view of the boxed region in **c**. The I61R/S67W/V101R triad (black spheres) remains juxtaposed to the pegRNA template and the extended NTS, recreating the stabilising clamp observed in the binary cryo-EM structure.

To confirm that the benefits translate to the full editor, we used AlphaFold 3 to predict a post-elongation assembly in which PEmaxΔRM3 remains docked beneath Cas9 and a nicked target DNA (Fig. 4c). The model recapitulates the same tripartite clamp on the pegRNA template (Fig. 4d), suggesting that the mutations stabilize the primer–template duplex throughout strand-displacement and gap-filling. We propose that this reinforced grip prolongs dwell time on the substrate, increases processivity, and ultimately boosts prime-editing efficiency in cells.

Together, these findings provide proof-of-concept for the application of saturated mutagenesis screens to optimize prime editors. We identified and validated three RT variants (I61R, V101R, S67W) that significantly enhance prime editing efficiency, particularly for small deletions. These variants synergize with both PEmax and epegRNAs, demonstrating their broad utility across different editing contexts and architectures. This work not only advances our understanding of RT engineering but also provides a robust framework for further optimizing prime editing systems for therapeutic and research applications.

## Discussion

In this study, we developed and optimized a compact prime editor, PE2ΔR, by removing the RNase H domain of M-MLV reverse transcriptase (RT). This modification not only reduced the size of the RT by 26.4% but also significantly enhanced prime editing efficiency across multiple genomic loci and edit types. To further enhance prime editing efficiency, we conducted a high-throughput saturated mutagenesis screen targeting two key DNA-binding regions within the fingers domain of RT (S60-Q84 and N95-D124)^4^. This screen identified several novel RT variants, including the triple mutant PE2ΔR-I61R, V101R, S67W (PE2ΔRM3), which increased editing efficiency by up to 86% compared to the parental PE2 in the eGFP reporter cells. The synergistic effects of these mutations highlight the importance of optimizing RT-DNA:RNA interactions, thermostability, and processivity in improving prime editing outcomes. Notably, the benefits of these mutations were not limited to PE2ΔR but extended to PEmax, where the corresponding variant (PEmaxΔRM3) achieved a 90% improvement in editing efficiency in the eGFP reporter cells.

The broad utility of PEmaxΔRM3 was further validated across diverse editing types, including base substitutions, small insertions, and small deletions. While improvements in base substitutions and small insertions were moderate (12%-37%), the most striking enhancement was observed for small deletions, where editing efficiency increased by up to 87%. This suggests that the optimized RT variants may preferentially stabilize the DNA:RNA hybrid during the introduction of deletions, or that the optimized RT functions with more seamless coordination with the subsequent DNA damage repair process. Additionally, the compatibility of PEmaxΔRM3 with engineered pegRNAs (epegRNAs) further highlights its potential for integration into advanced prime editing systems.

Our findings have several important implications for the field of genome editing. First, they demonstrate the feasibility of using high-throughput mutagenesis screens to identify and optimize RT variants for enhanced prime editing efficiency. This approach can be extended to other domains of RT or even other components of the prime editing machinery. Second, the improved editing efficiency of PE2ΔRM3 and PEmaxΔRM3 highlights the importance of compact, efficient PE *for in vivo* delivery applications, where size constraints and editing efficiency are critical considerations. Finally, the compatibility of these variants with epegRNAs suggests that future optimizations could combine RT engineering with pegRNA design to achieve even greater editing efficiencies.

Despite these advancements, several questions remain. For instance, the structural mechanisms by which the I61R, V101R, and S67W mutations enhance RT activity and prime editing efficiency are not fully understood. Structural studies and molecular dynamics simulations could provide insights into how these mutations stabilize the PE-DNA:RNA complex and improve processivity. Additionally, the performance of these variants in primary cells and animal models remains to be evaluated, as cellular context and chromatin state may influence editing outcomes.

In conclusion, this study provides a robust framework for optimizing prime editing systems through RT engineering. By combining domain truncation, high-throughput mutagenesis screen, and functional validation, we identified novel RT variants that markedly enhance prime editing efficiency, particularly for small deletions. These findings not only advance our understanding of RT function but also pave the way for the development of next-generation prime editors with improved therapeutic potential.

## Method

### Cell Lines and Culture Conditions

HEK293T cells were derived from human embryonic kidney 293 cells expressing SV40 T-antigen (ATCC, ATCC® CRL-3216™). 293FT cell line (Thermo Fisher) is a fast-growing, highly transfectable clonal isolated from H3K293T cells. HEK293T and 293FT cells were cultured in Dulbecco’s Modified Eagle’s Medium (DMEM) supplemented with 10% fetal bovine serum (FBS) and 1% penicillin/streptomycin. All cells were maintained in a 5% CO_2_ atmosphere at 37°C.

### Plasmids

The pCMV-PE2ΔR plasmid was constructed via deleting RNaseH domain from RT of pCMV-PE2 (#132775, Addgene). To construct pCMV-PE2ΔRM3, the mutations I61R, V101R, S67W were introduced into the RT of pCMV-PE2ΔR. Similarly, the RNaseH domain was deleted from RT of pCMV-PEmax (#174820, Addgene) to generate pCMV-PEmaxΔR. In addition, the pCMV-PEmaxΔRM3 plasmid was constructed by incorporating the I61R, V101R, S67W mutations into the RT of pCMV- PEmaxΔR. The protein sequences of PE2ΔR, PE2ΔRM3, and PEmaxΔRM3 are provided on Supplementary sequences.

All pegRNAs/epegRNAs and nicking sgRNAs used in this study were synthesized and cloned into vector pUC-GW-Amp, with their expressions driven by U6 promoter. A full list of plasmids is provided in Supplementary Table 1. The sequences of pegRNA/epegRNAs and nicking sgRNAs used in this study are listed in Supplementary Table 2 and Supplementary Table 3, respectively.

### Construction of eGFP reporter cell line

To generate the eGFP reporter cell line, we first replaced the Cas9 coding sequence (CDS) in the lentiCas9-Blast (#52962, addgene) with mutated eGFP CDS, generating the lenti-eGFP(Stop)-Blast construct. HEK293T cells were then infected with lenti-eGFP(Stop)-Blast lentiviruse for 16 h, followed by selection with 30 ug/ml blasticidin for 72 h. Subsequently, 500-1000 cells were plated into a 10-cm dish and cultured until clear colonies formed. We picked 30 single colonies and measured eGFP mRNA levels in each cell line. The single colony-derived cell line with the highest eGFP expression was selected for subsequent studies. The sequence of mutated eGFP is provided in Supplementary Sequences.

### Cell transfection and DNA preparation

To perform prime editing in HEK293T cells, 1x10^5^ cells were plated per well in a 24-well plate. After 24 h, the cells were co-transfected with 350 ng of the prime editor plasmid, 150 ng of the pegRNA plasmid, and 50 ng of the nicking sgRNA plasmid using lipofectamine 3000 (Invitrogen) according to the manufacturer’s instructions. After 72 h, FACS analysis was conducted on HEK293T eGFP reporter cells. To access editing efficiency at endogenous genomic loci, genomic DNA was extracted using the DNeasy kit (QIAGEN) 72 h post-transfection.

### Lentivirus packaging and transduction

To package lentiviruses, 293FT cells were seeded 6 x 10^7^ cells per 10-cm dish and allowed to recover overnight. The next day, cells were transfected with packaging plasmids using lipofectamine 3000 (Invitrogen) according to the manufacturer’s instructions. At 16 h after transfection, the medium was replaced with fresh cultural medium. Forty-eight hours after medium change, the virus-containing medium was collected, filtered through a 0.45 µm cellulose acetate filter, and stored at -80°C. For transduction, HEK293T cells were resuspended in fresh culture medium with supplementation with 6 µg/ml polybrene and lentivirus-containing supernatant for 16 h, followed by selection with blasticidin or puromycin for the time indicated in the text.

### Next generation sequencing and analysis of editing outcomes

To evaluate prime editing efficiency, genomic regions of each locus were amplified from DNA samples and sequenced using the Illumina Novaseq platform. Primers used for genomic DNA PCR amplification are listed in Supplementary Table 4. Sequencing reads were demultiplexed, and amplicon sequences were aligned to a reference sequence using CRISPResso2^23^. CRISPResso2 was run in standard mode with the ‘discard_indel_reads’ option enabled, while all other parameters were set to default. Editing efficiency was calculated as the percentage of non-discarded reads containing the desired edits without indels, divided by the total number of reads. Indel rate was determined as the proportion of discarded reads relative to the total number of reads. For all experiments, sequencing reads were analyzed for indels within a 30-nt window upstream and downstream of each pegRNA or sgRNA nick site, inclusive.

### Construction of saturated mutagenesis library

The construction of a mutagenesis library containing all possible mutations for S60-Q84 and N95-D124 (Supplementary Sequence 4) within the fingers domain of MMLV-RT was conducted following a previously described protocol with modifications^24^ . Briefly, we first inserted DNA sequence encoding prime editor PE2 lacking RNaseH domain and its promoter into lentiGuide-Puro vector (#52963, addgene). The oligos encoding all mutants (Supplementary Table 5), along with the wild-type sequence, were synthesized using a 12k oligo array (GenScript) and cleaved from the array as a single master oligo pool. Subsequently, these two individual libraries of mutants were PCR recovered from the master oligo pool using corresponding primer pairs. The PCR mixture included 10ng of chip oligos, 2 µM of 5’ primer, 2 µM of 3’ primer, and Phanta Max Master Mix (Vazyme) in a total reaction volume of 50 µl. PCR products were gel purified (Qiagen) and fused with the remaining RT sequence via Gibson Assembly (NEB). The assembled full-length of RT was then ligated into a linearized vector backbone using T4 ligase. The ligation products were electroporated into 25 µl of Electro MAX Stbl4 competent cells according to manufacture’s instructions (Life Technologies). Recovered bacterial cultures were plated onto 24-cm square LB-agarose plates containing ampicillin and incubated at 30°C for 48-72 h. Bacterial colonies were scrapped from the plates using ice-cold LB medium followed by extraction of library plasmids via maxi-preps.

To evaluate the coverage efficiency of each library, primers with Illumina adaptor were used to amplify the mutation regions. PCR products were gel-purified and sequenced on the NovaSeq 6000 platform (Novogene). The reads were analysed to quantify the representation of each mutant oligo in its library.

### Infection optimization for screening

Optimal infection conditions for each batch of virus were determined to achieve an infection efficiency of 20-40%, ensuring that most cells received only one stably integrated mutant. Infections were performed in a 12-well plate format, with 3x 10^6^ cells seeded in each well. To determine optimal virus volumes, cells were spin-infected (1,000 x g for 2 h at 33°C) with varying virus amounts (0, 50, 100, 200, 400, 600, 800, and 1000 µl) in the presence of 8 µg/ml polybrene. After spin-infection, the plate was returned to the incubator for a recovery of 24 h. After recovery, cells were trypsinized and replated in equal numbers into 2 wells of a 6-well plate containing complete medium, with one well supplemented with 0.5 µg/ml puromycin. Five days after selection, cell counts were compared between puromycin-selected and non-selected wells to calculate infection efficiency. Viral volumes yielding 20-40% infection efficiency were used for subsequent screening.

### Library screening and analysis

Screening was performed using the optimized viral volumes in a 12-well plate format, as described for viral titration. For libraries 1 and 2, 5x10^6^ and 6x10^6^ cells, respectively, were used to achieve a representation of 2000 cells per mutant following puromycin selection. Cells were selected with puromycin for 9 days, after which pegRNA and nicking RNA were transfected to enable prime editing, thereby restoring eGFP fluorescence. Two days after transfection, cells were trypsinized, resuspended in PBS, and subjected to FACS. eGFP-positive or eGFP-negative cells were collected separately for DNA extraction and next-generation sequencing as described above. For analysis, the log2 fold-change of each mutant in the eGFP-positive group was calculated relative to the eGFP-negative group for each biological replicate.

### Validation of screen hits

To validate RT mutants identified from the screen, each mutation was individually constructed or combined as needed in pCMV-PE2ΔR. Validation was firstly performed using the eGFP reporter cells in an arrayed format. For each condition, the prime editor plasmid harbouring the specific mutation(s) was co-transfected with the corresponding pegRNA and nicking sgRNA that allows for measurement of their efficiency in restoring eGFP expression. The PEMaxΔRM3 that contains I61R, V101R, S67W was constructed and used to evaluate its editing efficiency at endogenous genomic loci. The sequences for pegRNA/nicking sgRNA pairs used in this study were provided in Supplementary Table 2 and 3.

### Quantification and statistical analysis

All statistical analyses were performed using GraphPad Prism 10.4.1. Data were expressed as the mean ± standard deviation using at least three biologically independent replicates. The statistical differences between the two experimental groups were calculated using a heteroscedastic two-tailed student’s t-test. p < 0.05 was considered significant.

## Acknowledgements

These studies were supported by the National University of Singapore Start-up grant (H.J.Y.) and MOE Tier1 grant (NUHSRO/2021/114/T1/Seed-Sep/07, H.J.Y.).

## Author Contributions

Conceptualization, H. Y.;

Methodology, Y. W., C. Z. and H. Y.;

Formal Analysis, Y. W., C. Z., T. T., and H. Y.;

Investigation, Y. W., C. Z., T. T., Y. S., K. C., and H. Y.;

Writing-Original Draft, Y. W., C. Z. and H. Y.;

Supervision and Funding Acquisition, H. Y.

---

**Supplementary Figure 1.**
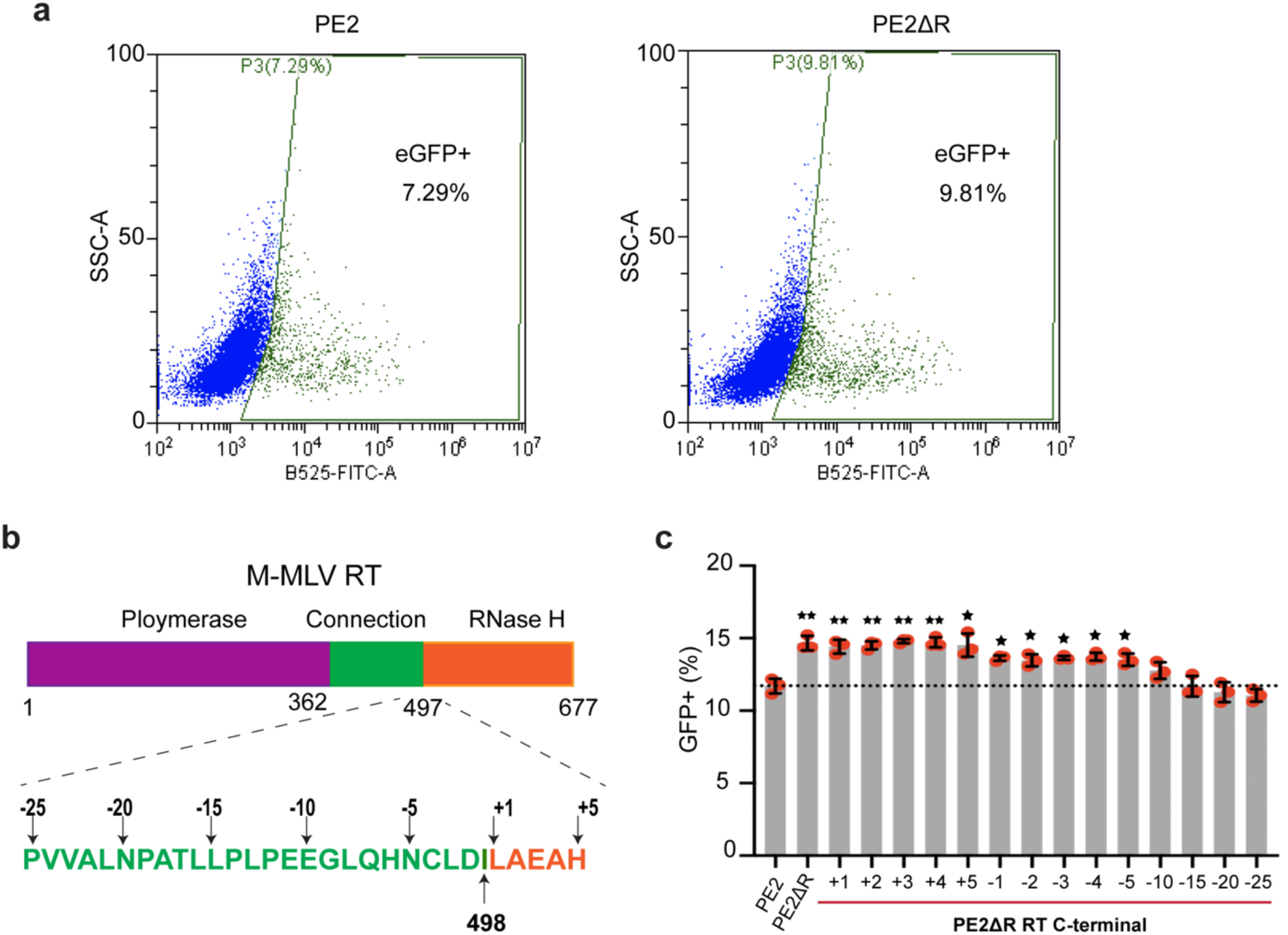
PE2ΔR overperforms PE2 in the eGFP reporter cell line. **a.** Representative flow cytometry analysis of eGFP+ cells upon prime editing via PE2 and PE2ΔR with using a nicking sgRNA. **b.** Schematic representation of the position of each RT truncation variant. **c.** Editing efficiency of PE2, PE2ΔR, and different PE2ΔR C-terminus truncation variants were quantified by flow cytometry. Results were obtained from three independent experiments. * p< 0.05, ** p< 0.01 by Student’s *t-*test. Error bar in c indicate mean ± SD.

**Supplementary Figure 2.**
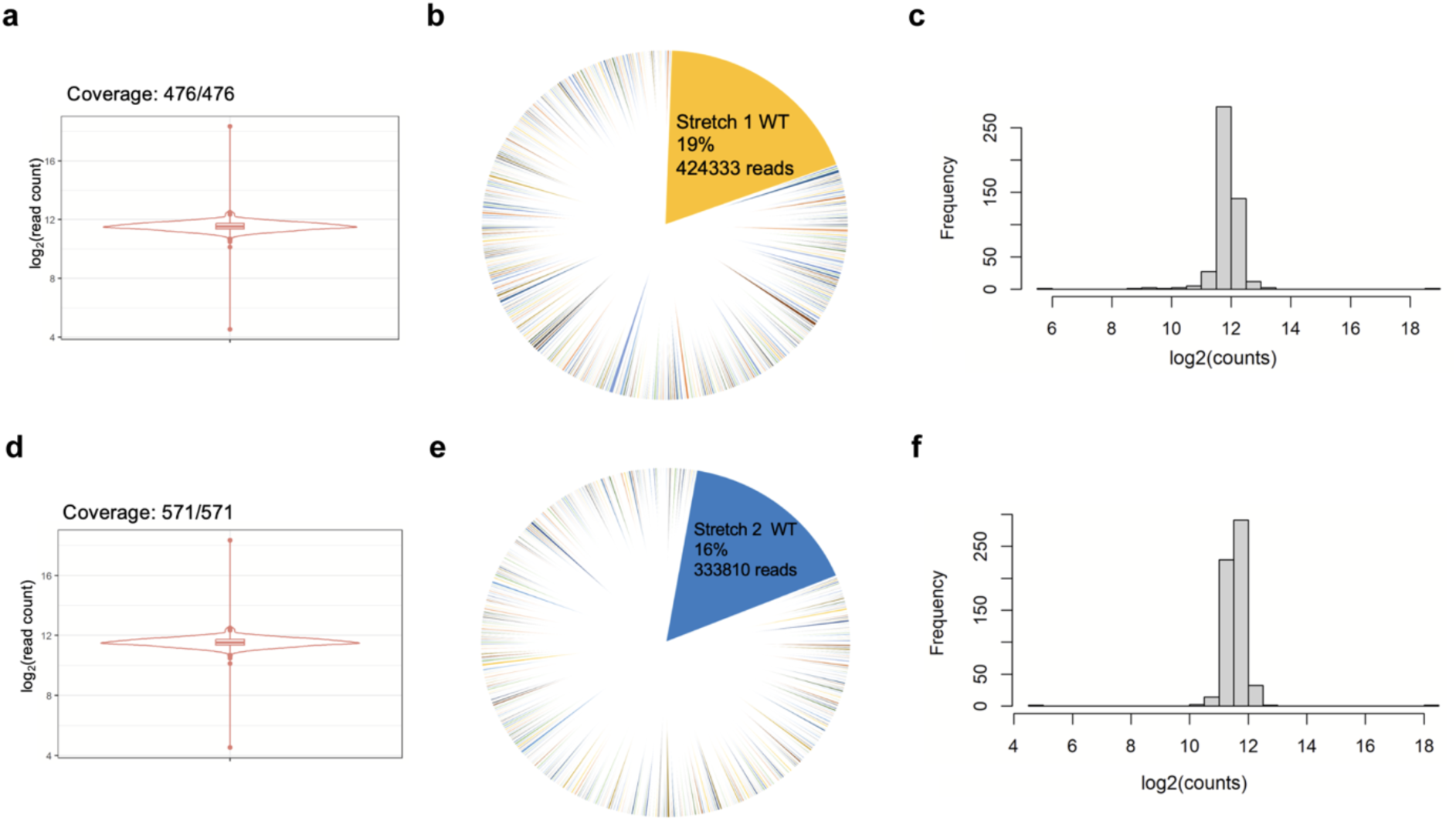
Deep sequencing analysis of the constructed saturated mutagenesis libraries. Violin plot **(a)**, pie chart **(b**) and histogram **(c)** showing the distribution of library members of the Stretch 1 mutant library. Violin plot **(d)**, pie chart **(e**) and histogram **(f)** showing the distribution of library members of the Stretch 2 mutant library. Each dot in the violin plot represents a singular member of the mutant library, and the frequency in the histogram refers to the number of members of the mutant library. The coverage of the library is indicated at the top of the violin plot. The wild-type (WT) proportion is indicated on the pie chart, along with the total number of sequencing reads of the wild-type.

**Supplementary Figure 3.**
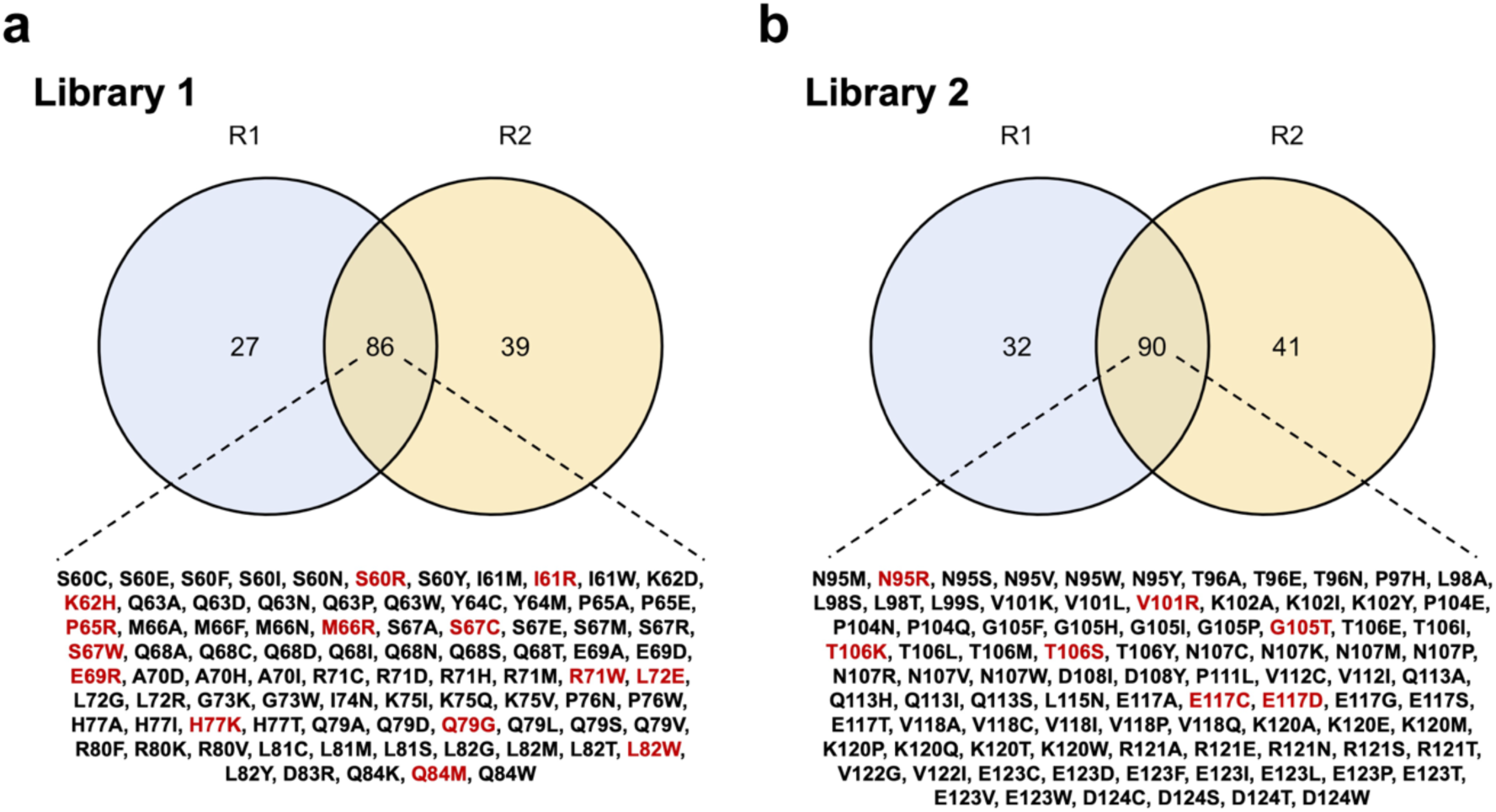
Candidate hits identified from mutagenesis screening. Venn Diagram illustrating the list of variants identified in both replicates of the screening. Variants selected for functional validation are highlighted in red.

**Supplementary Figure 4.**
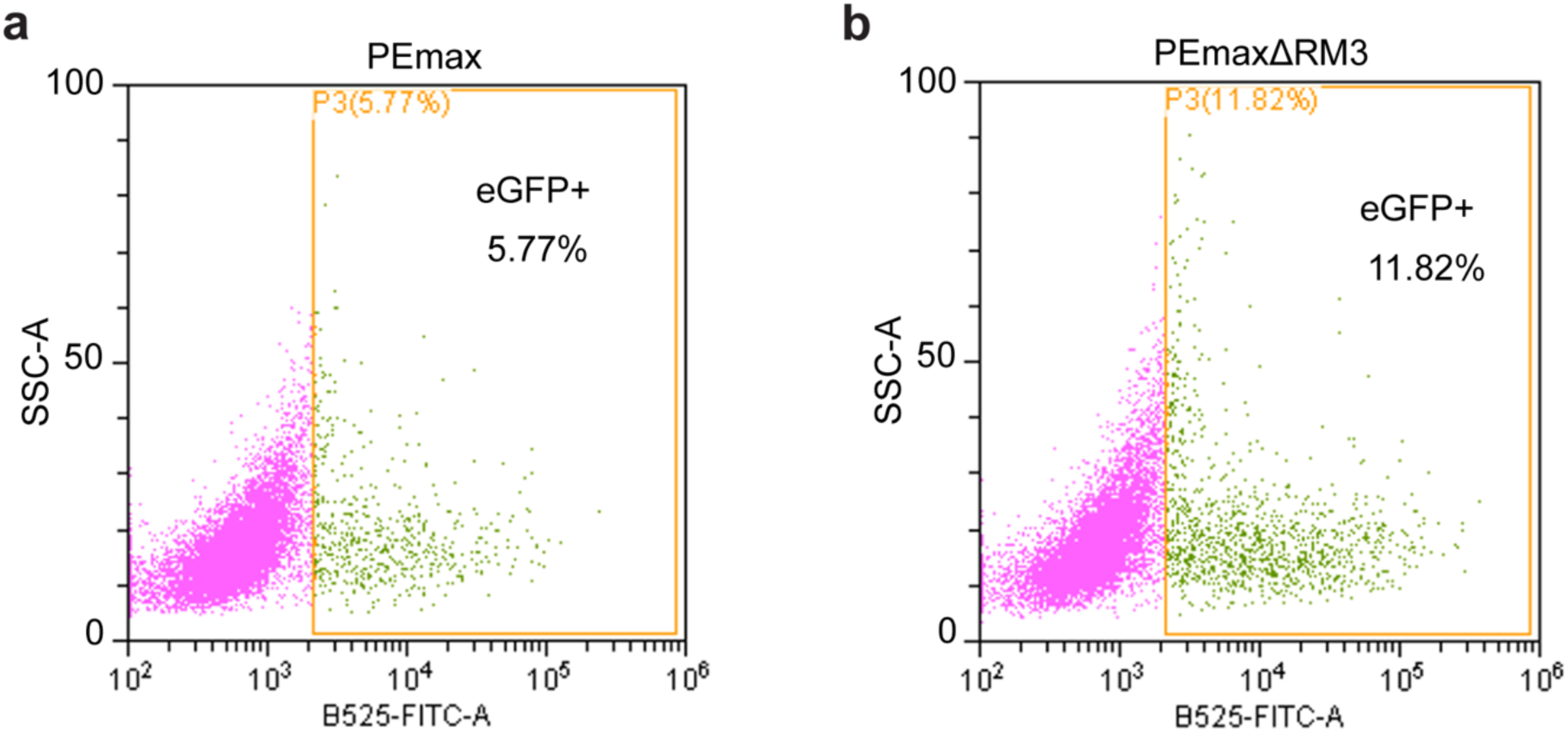
PEmaxΔRM3 overperforms PEmax in the eGFP reporter cell line. **a.** Representative flow cytometry analysis of eGFP+ cells upon prime editing via PEmax and PEmaxΔRM3.

## Reference

1 Anzalone, A. V. et al. Search-and-replace genome editing without double-strand breaks or donor DNA. Nature 576, 149–157 (2019). 10.1038/s41586-019-1711-4

2 Chen, P. J. & Liu, D. R. Prime editing for precise and highly versatile genome manipulation. Nat Rev Genet 24, 161–177 (2023). 10.1038/s41576-022-00541-1

3 Zhao, Z., Shang, P., Mohanraju, P. & Geijsen, N. Prime editing: advances and therapeutic applications. Trends Biotechnol 41, 1000–1012 (2023). 10.1016/j.tibtech.2023.03.004

4 Oscorbin, I. P. & Filipenko, M. L. M-MuLV reverse transcriptase: Selected properties and improved mutants. Comput Struct Biotechnol J 19, 6315–6327 (2021). 10.1016/j.csbj.2021.11.030

5 Velimirovic, M. et al. Peptide fusion improves prime editing efficiency. Nat Commun 13, 3512 (2022). 10.1038/s41467-022-31270-y

6 Doman, J. L., et al. Phage-assisted evolution and protein engineering yield compact, efficient prime editors. Cell 186, 3983–4002 e3926 (2023). 10.1016/j.cell.2023.07.039

7 Chen, P. J. et al. Enhanced prime editing systems by manipulating cellular determinants of editing outcomes. Cell 184, 5635–5652 e5629 (2021). 10.1016/j.cell.2021.09.018

8 Lan, T., et al. Mini-PE, a prime editor with compact Cas9 and truncated reverse transcriptase. Mol Ther Nucleic Acids 33, 890–897 (2023). 10.1016/j.omtn.2023.08.018

9 Weber, Y., et al. Enhancing prime editor activity by directed protein evolution in yeast. Nat Commun 15, 2092 (2024). 10.1038/s41467-024-46107-z

10 Yan, J. et al. Improving prime editing with an endogenous small RNA-binding protein. Nature 628, 639–647 (2024). 10.1038/s41586-024-07259-6

11 Gao, Z. et al. A truncated reverse transcriptase enhances prime editing by split AAV vectors. Mol Ther 30, 2942–2951 (2022). 10.1016/j.ymthe.2022.07.001

12 Grunewald, J., et al. Engineered CRISPR prime editors with compact, untethered reverse transcriptases. Nat Biotechnol 41, 337–343 (2023). 10.1038/s41587-022-01473-1

13 Bock, D. et al. In vivo prime editing of a metabolic liver disease in mice. Sci Transl Med 14, eabl9238 (2022). 10.1126/scitranslmed.abl9238

14 Zheng, C., et al. A flexible split prime editor using truncated reverse transcriptase improves dual-AAV delivery in mouse liver. Mol Ther 30, 1343–1351 (2022). 10.1016/j.ymthe.2022.01.005

15 Liu, P., et al. Improved prime editors enable pathogenic allele correction and cancer modelling in adult mice. Nat Commun 12, 2121 (2021). 10.1038/s41467-021-22295-w

16 Zhang, W. et al. Enhancing CRISPR prime editing by reducing misfolded pegRNA interactions. Elife 12 (2024). 10.7554/eLife.90948

17 Nelson, J. W. et al. Engineered pegRNAs improve prime editing efficiency. Nat Biotechnol 40, 402–410 (2022). 10.1038/s41587-021-01039-7

18 Baranauskas, A. et al. Generation and characterization of new highly thermostable and processive M-MuLV reverse transcriptase variants. Protein Eng Des Sel 25, 657–668 (2012). 10.1093/protein/gzs034

19 Lin, Q. et al. High-efficiency prime editing with optimized, paired pegRNAs in plants. Nat Biotechnol 39, 923–927 (2021). 10.1038/s41587-021-00868-w

20 Kim, H. K. et al. Predicting the efficiency of prime editing guide RNAs in human cells. Nat Biotechnol 39, 198–206 (2021). 10.1038/s41587-020-0677-y

21 Kim, H. K. et al. SpCas9 activity prediction by DeepSpCas9, a deep learning-based model with high generalization performance. Sci Adv 5, eaax9249 (2019). 10.1126/sciadv.aax9249

22 Najmudin, S. et al. Crystal structures of an N-terminal fragment from Moloney murine leukemia virus reverse transcriptase complexed with nucleic acid: functional implications for template-primer binding to the fingers domain. J Mol Biol 296, 613–632 (2000). 10.1006/jmbi.1999.3477

23 Clement, K. et al. CRISPResso2 provides accurate and rapid genome editing sequence analysis. Nat Biotechnol 37, 224–226 (2019). 10.1038/s41587-019-0032-3

24 Read, A., Gao, S., Batchelor, E. & Luo, J. Flexible CRISPR library construction using parallel oligonucleotide retrieval. Nucleic Acids Res 45, e101 (2017). 10.1093/nar/gkx181

